# *ZNF251* haploinsufficiency confers PARP inhibitors resistance in *BRCA1*-mutated cancer cells through activation of homologous recombination

**DOI:** 10.1101/2022.09.29.510119

**Authors:** Huan Li, Srinivas Chatla, Xiaolei Liu, Zhen Tian, Umeshkumar Vekariya, Peng Wang, Dongwook Kim, Stacia Octaviani, Zhaorui Lian, George Morton, Zijie Feng, Dan Yang, Katherine Sullivan-Reed, Wayne Childers, Xiang Yu, Kumaraswamy Naidu Chitrala, Jozef Madzo, Tomasz Skorski, Jian Huang

**Author notes:** Correspondence: Jian Huang,; Tomasz Skorski. Surgical Research Lab, Department of Surgery, Cooper University Health Care, Camden, NJ, United States. Department of Engineering Technology, University of Houston, Houston, Texas, United States (Kumaraswamy Naidu Chitrala). These authors contributed equally to this work (Huan Li, Srinivas Chatla). These authors contributed equally to this work (Xiaolei Liu, Zhen Tian).

## Abstract

Poly (ADP-ribose) polymerase inhibitors (PARPis) represent a promising new class of agents that have demonstrated efficacy in treating various cancers, particularly those with *BRCA1/2* mutations. Cancer-associated *BRCA1/2* mutations disrupt DNA double-strand break (DSB) repair by homologous recombination (HR). PARP inhibitors (PARPis) have been used to trigger synthetic lethality in *BRCA1/2*-mutated cancer cells by promoting the accumulation of toxic DSBs. Unfortunately, resistance to PARPis is common and can occur through multiple mechanisms, including the restoration of HR and/or stabilization of replication forks. To gain a better understanding of the mechanisms underlying PARPis resistance, we conducted an unbiased CRISPR-pooled genome-wide library screen to identify new genes whose deficiency confers resistance to the PARPi olaparib. Our research revealed that haploinsufficiency of the *ZNF251* gene, which encodes zinc finger protein 251, is associated with resistance to PARPis in various breast and ovarian cancer cell lines carrying *BRCA1* mutations. Mechanistically, we discovered that *ZNF251* haploinsufficiency leads to stimulation of RAD51-mediated HR repair of DSBs in olaparib-treated *BRCA1*-mutated cancer cells. Moreover, we demonstrated that a RAD51 inhibitor reversed PARPi resistance in *ZNF251* haploinsufficient cancer cells harboring *BRCA1* mutations. Our findings provide important insights into the mechanisms underlying PARPis resistance by highlighting the role of RAD51 in this phenomenon.

## Introduction

Poly (ADP-ribose) polymerases (PARPs), also known as NAD+ ADP-ribosyltransferases, are an emerging family of 18 enzymes that can catalyze the transfer of ADP-ribose to target proteins (poly ADPribosylation)(1,2). PARPs play an important role in various cellular processes, including the modulation of chromatin structure, transcription, replication, recombination, and DNA repair(3). PARP1 is the most potent enzyme in this group, accounting for 80–90% of DNA damage-induced PARylation, and plays a key role in the DNA damage response (DDR), including the repair of DNA single-strand breaks (SSBs) and double-strand breaks (DSBs)(4-6). SSBs are repaired using PARP1-mediated base excision repair (BER). DSBs can be repaired by three classical pathways: *BRCA1/2*-dependent homologous recombination (HR), DNA-PKC-mediated non-homologous end joining (NHEJ), and PARP1-mediated alternative NHEJ (Alt-NHEJ). These DNA repair pathways can either work independently or in coordination to prevent or repair different types of DSBs.

Mutations in *BRCA1/2* genes, leading to dysfunctional HR, pose a significant risk for the development of breast and ovarian cancers. BRCA1 and BRCA2 interact with various proteins involved in the HR repair pathway and are essential for this process, operating at different stages of DSB repair. Since PARP inhibitors (PARPis) induce DSBs in cells with defective HR, cells harboring *BRCA1/2* mutations are highly susceptible to PARP inhibitor treatments(3-6). Currently, FDA-approved PARPis, including olaparib, rucaparib, niraparib, and talazoparib, act as competitors to NAD+, thereby inhibiting poly (ADP ribose) polymerase activity. PARP1 inhibition leads to the accumulation of single-strand breaks (SSBs), which results in toxic DSBs and synthetic lethality in the absence of functional HR. Unfortunately, the majority of patients with *BRCA1/2* mutated tumors initially respond positively to PARPis treatment but eventually develop resistance, leading to disease relapse and progression.

To investigate the mechanisms underlying resistance to PARP inhibitors (PARPis), we conducted a genome-wide CRISPR screen to identify the gene mutations that confer resistance to olaparib. Our findings revealed that haploinsufficiency of the zinc finger 251 (*ZNF251*) gene, resulting in partial knockdown of the ZNF251 protein (referred to as *ZNF251*KD), causes resistance to olaparib in multiple *BRCA1*-mutated cancer cell lines. Moreover, we observed that breast cancer cells with *BRCA1* mutation and *ZNF251*KD (*BRCA1*mut+*ZNF251*KD) were not only resistant to various PARPis but also to platinum-based drugs and DNA polymerase theta (Polθ) inhibitors.

Our study further demonstrated that activation of HR and replication fork stabilization is associated with resistance to PARPis conferred by *ZNF251*KD in *BRCA1*-mutated cells. Critically, we also showed that *BRCA1*mut+*ZNF251*KD breast cancer cells were sensitive to RAD51 inhibitor (RAD51i), which also restored their sensitivity to PARPi. These results suggest that *ZNF251*KD-mediated resistance to PARPis involves the HR pathway, which may represent a therapeutic target for overcoming PARPis resistance in *BRCA1*mut+*ZNF251*KD breast cancer cells.

## Materials and Methods

### Cell lines and cell culture

MDA-MB-436 and HCC1937 cells were purchased from ATCC. MDA-MB-436 and Ovcar8 cells were maintained in DMEM supplemented with 10% fetal bovine serum (FBS) and 1% penicillin/streptomycin. HCC1937 cells were cultured in RPMI 1640 medium supplemented with 10% FBS and 1% penicillin/streptomycin. All cell lines were analyzed and authenticated by morphological inspection and biochemical examination of the *BRCA1* mutation pathway as well as short tandem repeat profiling analysis. Mycoplasma testing was performed to exclude the possibility of mycoplasma contamination in all cell lines.

### Chemical Compounds

Olaparib (Catalog# A4154), cisplatin (Catalog# A8321), carboplatin (Catalog# A2171), and 5-fluorouracil (Catalog# A4071) were purchased from APExBIO. UPF 1096 (Catalog# S8038), NMS-P118(Catalog# S8363), stenoparib (E7449) (Catalog# S8419), niraparib (Catalog# S2741), rucaparib (Catalog# S4948), and veliparib (ABT-888) (Catalog# S1004) were purchased from Selleckchem. Cisplatin (Catalog# A10221) was purchased from AdooQ Bioscience. ART-558(Catalog# HY-141520), RAD51 inhibitor RI-1 (Catalog# HY-15317), and olaparib (for *in vivo* experiment: Catalog# HY-10162) were purchased from MedChem Express (MCE). ART-812 was synthesized at the Moulder Center for Drug Discovery, Temple University School of Pharmacy. All compounds were dissolved, aliquoted, and stored in accordance with the manufacturer’s instructions.

### Pooled Genome-wide CRISPR/Cas9 Screen

The GeCKO CRISPR library was purchased from Addgene (#1000000048), amplified, and packaged as a lentivirus based on the instructions on the Addgene website. CRISPR screening was performed as described previously(7). Briefly, MDA-MB-436 cells were transduced with lentivirus carrying the GeCKO library, and puromycin selection was performed for 3 days. The transduced MDA-MB-436 cells were treated with olaparib for 14 days. The medium was changed by adding fresh olaparib every three days during 14 days screen, and the surviving cells were harvested. Genomic DNA was extracted, and PCR was performed before deep sequencing of the sgRNA sequence in the genome of surviving cells. All deep sequencing data are available from GEO (series accession number GSE205221). For data analysis, we calculated the enrichment score as follows: enrichment score= (sgRNA number from the reads)/ (sgRNA number in the library) × log_2_ (average abundance). sgRNAs used for validation were synthesized and constructed as previously described(7). Primer sequences are listed in Supplementary Table S1.

### T7EN1 assays and DNA sequencing

The T7EN1 assay was performed as previously described (7). To identify *ZNF251* mutations, the purified PCR product was cloned into the pCR2.1-TOPO TA vector (TOPO TA cloning kit; Life Technologies) and sequenced by Sanger sequencing. Primers used for Sanger sequencing are listed in Supplementary Table S1.

### Generation of mutant single clones

Five hundred transduced MDA-MB-436 cells were mixed with 1 ml of methylcellulose (MethoCult H4034 Optimum, Stem Cell Technologies) in a 6-well cell culture plate and cultured at 37 °C in a 5% CO_2_ incubator. Two weeks later, single colonies were selected and cultured in a 96-well plate with complete medium supplemented with 2% penicillin/streptomycin. The cells were passaged every two or three days, and 1/3 of the cells were collected for genomic DNA extraction. The *ZNF251* target region was PCR-amplified and sequenced.

### Cell viability assay

Cells (1×10^4^) were cultured in 100μ μL of complete medium in a 96-well plate and treated as indicated. Cell viability was measured at different time points as described in the trypan blue exclusion viability test. The final number of viable cells was calculated based on a standard growth curve. All key viability experiments were confirmed by the MTS assay (Promega, Catalog# G3582) and CCK-8 assay (APExBIO company, Catalog# K1018).

### Off-target effect examination

Off-target sites were predicted using an online search tool (http://crispr.mit.edu). 3bp mismatches compared to the target consensus sequence were allowed. The predicted off-target sequences were searched using the UCSC browse, and 500bp flanking the sites were PCR-amplified in primary cells and single mutation clones. The PCR product was subjected to a T7EN1 assay to determine the mutation. The PCR product was cloned into a TA vector and Sanger sequenced to identify the mutations.

### *ZNF251* complementation experiment

Exponentially growing MDA-MB-436 *ZNF251* WT and KD cells were seeded in six-well plates (1 million cells/well) and transfected with pcDNA3.1 vector or human *ZNF251* on pcDNA3.1 plasmid carrying the neomycin resistance (neo) gene. After transfection with 1μg and 2 μg plasmids, the cells were selected with G418 (400 μg/ml) in the culture medium for 2 weeks to maintain the selection of neomycin-resistant cells to generate stably transfected cell lines(8,9). Cells were plated in 96-well plates at a density of 1× 10^4^ cells/well in triplicate. The next day, the transfected cells were treated with DMSO or olaparib for three days, and cell viability was measured.

### Cell Viability, Apoptosis, and Proliferation Assays

Apoptosis and proliferation were analyzed using flow cytometry. Cells were stained using an FITC Annexin V apoptosis detection kit (BioLegend Catalog#641904) and a Brilliant Violet 423™ anti-Ki67 antibody (BioLegend Catalog#652411) following fixation and permeabilization. This process was performed 72 h after treatment with either 8μM Olaparib or an equivalent volume of DMSO as a control.

### Immunoblot analysis

Nuclear and total cell lysates were obtained as described previously (10) and analyzed by SDS-PAGE using primary antibodies against ATM (Santa Cruz Biotechnology #sc-135663), CtIP (Abcam #ab-70163), 53BP1 (Abcam #ab-175933), SLFN11 (Santa Cruz Biotechnology #sc-515071), BRCA1 (ThermoFisher Scientific #MA1-23164), BRCA2 (Santa Cruz Biotechnology #sc-28235), PALB2 (Proteintech #14340-1-AP), RAD51 (Abcam #ab-88572), RAD52 (Santa Cruz Biotechnology #sc-365341), RAD54 (Santa Cruz Biotechnology #sc-374598), and the following secondary antibodies conjugated to horseradish peroxidase (HRP): goat anti-rabbit (EMD Millipore #12–348) and goat anti-mouse (EMD Millipore #AP181P). ZNF251 western blot analysis was performed using ZNF251 antibody (Proteintech cat# 25601-1-AP) and GAPDH antibody (Cell Signaling Technology, cat#2118). For quantification of the western blot analysis, ImageJ software was used to measure the density of the protein bands.

### DNA damage/repair assays

DSBs were detected using a neutral comet assay, as described previously (10) with modifications. Briefly, comet assays were performed using the Oxiselect Comet Assay Kit (Cell Biolabs #STA-355), according to the manufacturer’s instructions. Images were acquired using an inverted Olympus IX70 fluorescence microscope with an FITC filter, and the percentage of tail DNA in individual cells was calculated using the OpenComet plugin of ImageJ. HR was measured using the DR-GFP reporter cassette as described previously (10). Briefly, the reporter plasmid was digested with I-SceI endonuclease, and the repaired GFP cells were counted by flow cytometry. The result was calculated as the total number of restored GFP-positive cells/total transfected M-cherry- or BFP-positive cells.

### Mice and *in vivo* studies

6-8 weeks-old female NOD/SCID/IL-2Rγ (NSG) mice (Jackson Laboratories) were subcutaneously injected with 1x10^6^ MBA-MD-436 cells in the flank. Mice were randomized into treatment groups when the tumor sizes reached 50-60 mm^3^. All animals with wild-type or *ZNF251*KD tumors of 50-60 mm^3^ were randomized into two groups (n=4/group), which were intraperitoneally administered vehicle or olaparib (10 mg/kg) daily for four weeks, respectively. From the start of the experiment, tumor volumes (V) were measured every three days based on the formula V = L×W^2^×0.5, where L represents the largest tumor diameter and W represents the perpendicular tumor diameter(11). After four weeks, all mice were euthanized, and tumors were dissected, imaged, weighed, or used for further characterization. All experiments involving animals were approved by the Institutional Animal Care and Use (IACUC) Committee of Cooper University.

### Fork protection assay/DNA fiber assay

At stalled forks, the degradation of DNA fibers was assessed as follows. Exponentially growing MDA-MB 436 *ZNF251* WT and KD cells were treated with 5 μM olaparib for 48 hours. Cells were sequentially pulse-labeled with 50 μM 5-chloro-2′-deoxyuridine thymidine (CldU) (Sigma-Aldrich) and 250 μM idoxuridine (IdU) (Sigma-Aldrich) for 30 min each, washed once with 1× PBS, and treated with 4 mM HU for 4 h. The cells were collected and resuspended in 1× PBS at a concentration of 500 cells/μL. 2.5 μl of the cell suspension was diluted with 7.5 μl of lysis buffer (200 mM Tris-HCl pH 7.5, 50 mM EDTA, and 0.5% [w/v] SDS) on a glass slide and incubated for 8 min at RT. The slides were tilted at 15°–60°, air-dried, and fixed with 3:1 methanol/acetic acid for 10 min. The slides were denatured with 2.5 M HCl for 90 min, washed with 1× PBS, and blocked with 2% BSA (Carl Roth) in PBS for 40 min. The newly replicated CldU and IdU tracks were labeled for 1.5 hr with anti-BrdU antibodies recognizing CldU (1:300, Abcam) and IdU (1:100, BD Biosciences), followed by 1 h incubation with anti-mouse Alexa Fluor 594 (1:500, #A11062, Life Technologies) and anti-rat Alexa Fluor 488 (1:500, #A21470, Life Technologies) secondary antibodies. Incubations were performed in the dark in a humidified chamber. After five washes in PBST for 3 min, the coverslips were mounted with 20 μL mounting media. DNA fibers were visualized using a Leica SP8 Confocal microscope at 63X objective magnification and images were analyzed using ImageJ software.

### Bioinformatics analysis of *ZNF251* expression in the cells sensitive and resistant to PARPi olaparib

To analyze the expression of *ZNF251* in cells sensitive and resistant to the PARPi olaparib, we performed bioinformatics analysis. Datasets for the respective inhibitors were downloaded from the Gene Expression Omnibus database (http://www.ncbi.nlm.nih.gov/geo), a large public repository for high-throughput molecular abundance data, specifically gene expression data(12). Dataset GSE165914 was used for the analysis of *ZNF251* expression in olaparib-sensitive and-resistant cells(13). Statistical analyses were performed using the GraphPad Prism 9.

### Quantification and Statistical Analysis

All statistical analyses were performed using the GraphPad Prism version 8. Cell viability data were analyzed using two-way ANOVA. Neural comet assay data were analyzed using the Mann-Whitney Rank Sum Test. Data from the DNA repair assay and *in vivo* experiments were analyzed using an unpaired t-test with Welch’s correction.

## Data availability

The data generated in this study are available in the article and its supplementary files. All deep sequencing data from our CRISPR screen are available at GEO (series accession number: GSE205221).

## Results

### A genome-wide CRISPR screen identified ZNF251 as a critical factor regulating sensitivity of BRCA1-mutated cells to PARPis

To identify genes whose deficiency confers drug resistance to the PARPi olaparib, we performed a genome-wide CRISPR genetic screen in MDA-MB-436 cells, a human breast cancer line harboring a *BRCA1* 5396 + 1G>A mutation in the splice donor site of exon 20, resulting in a BRCT domain-truncated protein. MDA-MB-436 cells are known to exhibit sensitivity to PARPis(14). We used the GeCKO CRISPR library, which has been demonstrated to be an efficient tool to screen for mutations that confer resistance to a BRAF inhibitor in a melanoma line(15). First, we packed the library into lentivirus with an optimal titer at a multiplicity of infection (MOI) of 0.3-0.4 and transduced MDA-MB-436 cells. After viral transduction, MDA-MB-436 breast cancer cells with either 0.3 μM or 1μM olaparib, an optimal dose chosen based on our preliminary tests (Supplementary Fig. 1A). After 14 days of treatment, we harvested living cells from the olaparib-treated group and extracted genomic DNA for polymerase chain reaction (PCR) of the region containing sgRNAs. Next, we conducted next-generation sequencing (deep sequencing) to identify sgRNAs that were enriched in olaparib-resistant cells (Supplementary Fig. 1B). For several genes, we found enrichment of multiple sgRNAs, suggesting that deficiency in these genes contributes to olaparib resistance. We then ranked the positive hits by the number of sgRNAs and enrichment changes per sgRNA. Interestingly, we identified several zinc finger genes as the highest-ranking genes in our screen. Among them, *ZNF251* and *ZNF518B* were the only two top hits recovered on the screen at the two doses (Supplementary Fig. 1C). Importantly, our screen identified several known genes whose loss-of-function causes olaparib resistance. These include *TP53BP1*(16), components of the Shieldin complex (*C20orf196* and *FAM35A*)(17), *DYNLL1*(18,19), and *EZH2*(20). This result, as shown in Supplementary Fig. 1D, validates the effectiveness of our screening process. Next, we tested both *ZNF251* and *ZNF518B* and found that targeting *ZNF251* resulted in stronger resistance to olaparib. Therefore, we chose *ZNF251* for this study.

To further validate whether deficiency of *ZNF251* confers resistance to olaparib, we used three newly designed sgRNAs to disrupt *ZNF251* in MDA-MB-436 breast cancer cells. We transduced cells with lentivirus-carrying sgRNAs specifically for *ZNF251* and performed the T7 Endonuclease I assay five days after transduction to determine the disruption efficiency. The efficiency of gene disruption ranged from 52.3% to 89% for all sgRNAs tested (Fig. 1A, top panel). Next, we used these cells to test whether the disruption of *ZNF251* could confer resistance to olaparib. Consistent with our screening data, we found that *ZNF251-*deficient cells showed marked resistance to olaparib treatment compared to the parental cells (Fig. 1A, bottom panel). Because the CRISPR/Cas9 genome editing system can create a spectrum of insertions/deletions (in/dels) in a cell population, we also isolated three *ZNF251-*deficient single clones, TOPO cloned and sequenced the PCR product encompassing the targeted region of gRNAs. Approximately 50% of the clones contained Cas9-mediated mutations, including deletions and insertions, at or near the sgRNA PAM sequences (Supplementary Fig. 1E), indicating that the in/dels were all monoallelic mutations. While it is more common for CRISPR to generate biallelic mutations in a gene, it is not rare to generate monoallelic mutations. To further confirm the heterozygosity of the *ZNF251-*deficient clones, we performed western blot analysis to quantify ZNF251 protein levels in the cells. Our results showed a significant reduction of approximately 50% in protein levels compared to wild-type control cells (Fig. 1B, top panel). These findings provide strong evidence that *ZNF251-*deficient clones are heterozygous knockdowns. Throughout the manuscript, we have referred to the mutation caused by *ZNF251* haploinsufficiency as “*ZNF251* knockdown” (*ZNF251*KD) to accurately describe it.

Next, we tested the drug resistance of three independent *ZNF251*KD clones (#1-3) to olaparib. Consistent with the data from the heterogeneous population of CRISPR-mutated cells, all three *ZNF251*KD clones showed resistance to olaparib compared to the parental (*ZNF251*WT) cells (Fig. 1B bottom panel). The IC_50_ of *ZNF251*-knockdown clones to olaparib was between 7.04 and 16.03 μM, whereas the IC_50_ for parental cells was 4.36 μM. Importantly, transfecting *ZNF251*KD cells with an ectopic expression plasmid carrying wild-type *ZNF251* cDNA fully reversed their resistance to olaparib, indicating that *ZNF251* haploinsufficiency is responsible for this resistance (Supplementary Fig. 1F). Consistent with their resistance to olaparib, *ZNF251*KD breast cancer cells exhibited increased proliferation and reduced apoptosis after olaparib treatment (Supplementary Fig. 2A, B). To address the question of whether resistance is correlated with *BRCA1* mutation, we knocked down *ZNF251* in isogenic *BRCA1-*wildtype and mutated HCC1937 human breast cancer cell lines. We found that *ZNF251* knockdown caused olaparib resistance in *BRCA1*-mutated but not *BRCA1*-wildtype breast cancer cells (Supplementary Fig. 3A, B).

We subsequently assessed whether *ZNF251*KD causes PARPi resistance *in vivo*. We experimentally tested the effects of olaparib on the growth of parental (*ZNF251*WT) and *ZNF251*KD3 MDA-MB-436 cell xenografts in immunodeficient NSG mice (Fig. 1C). First, we injected either 1x10^6^ wildtype or *ZNF251*KD3 cells subcutaneously into the flanks of 16 NSG female mice (eight and eight mice injected with either *ZNF251*WT or *ZNF251*KD3 cells).

**Figure 1.**
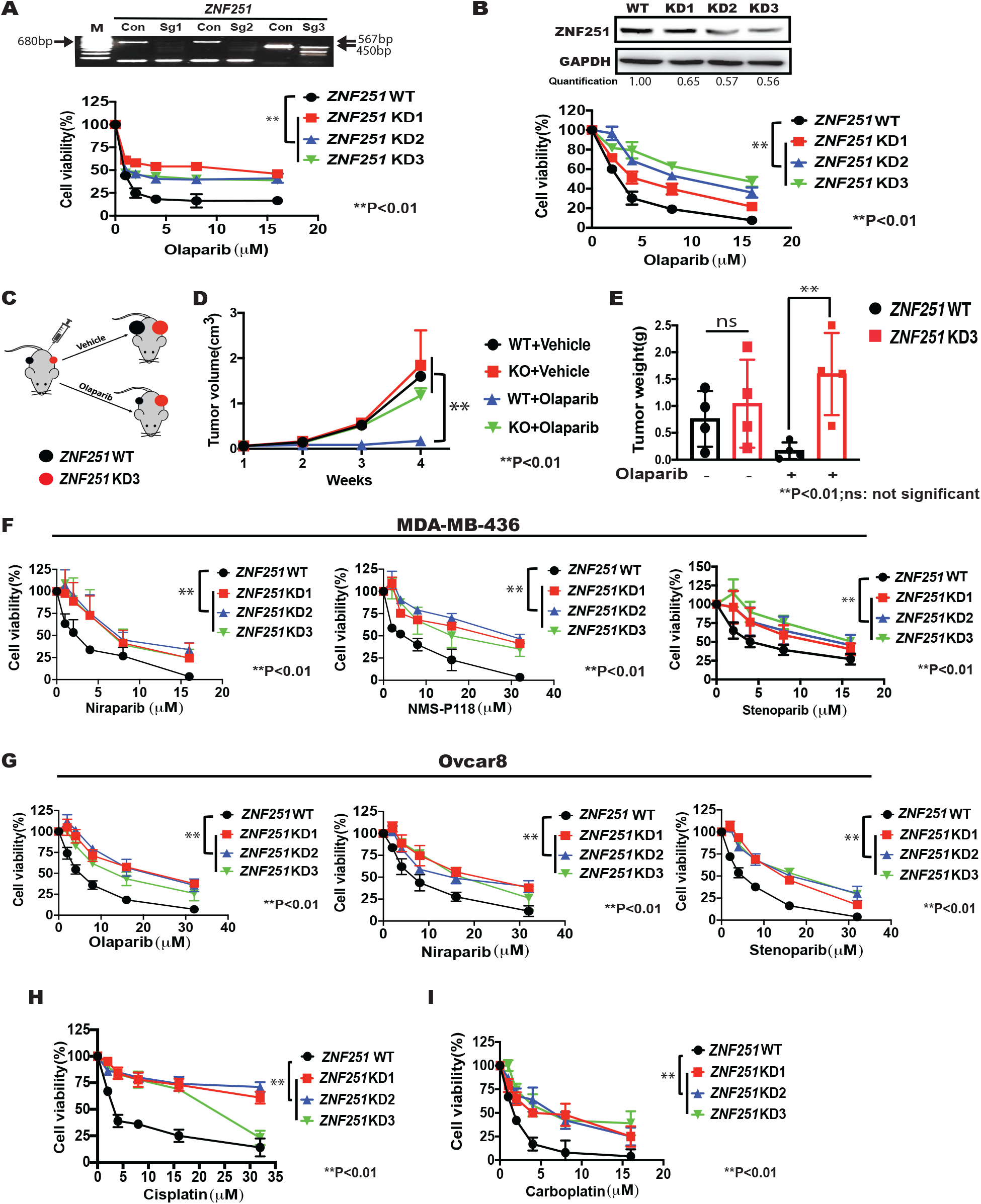
A genome-wide CRISPR screen in MDA-MB-436 *BRCA1*-mutated breast cancer cells uncovered *ZNF251* genes whose haploinsufficiency confers olaparib resistance. A. Top panel: T7EN1 assay analysis of specific sgRNA-mediated in/dels at *ZNF251* locus in MDA-MB-436 cells. Bottom panel: Cell growth curve of parental (*ZNF251* WT) and *ZNF251* KD MDA-MB-436 breast cancer cells following treatment with olaparib. The results represent three independent experiments. B. Top panel: Western blot analysis of *ZNF251*WT and *ZNF251*KD clones 1-3 of MDA-MB-436 breast cancer cells. GADPH was used as a loading control. The abundance of ZNF251 bands relative to the corresponding GADPH bands was assessed densitometrically. Bottom panel: Cell growth curve of parental (*ZNF251* WT) and *ZNF251* KD MDA-MB-436 breast cancer cells following treatment with olaparib. The results represent three independent experiments. C. Schematic of *in vivo* olaparib treatment experiment. D, E. The effect of olaparib on the growth of *ZNF251*WT and *ZNF251*KD MDA-MB-436 breast cancer cells xenografts in immune-deficient NOD/SCID/IL-2Rγ (NSG) mice was tested. The results represent three independent experiments. F, G. *ZNF251*KD caused multiple PARPis resistance in *BRCA1*-mutated breast (MDA-MB-436) and ovarian (Ovcar8) cancer lines. *ZNF251*KD was constructed in MDA-MB-436 and Ovcar8 cell lines and the resistance to olaparib, niraparib, NMS-P118, stenoparib was measured. H, I. The resistance of *ZNF251*WT and *ZNF251*KDs to cisplatin (H), carboplatin (I) was tested. The results represent three independent experiments.

Notably, tumors were observed in all 16 animals transplanted with MDA-MB-436 cells for ∼3-4 weeks. Next, all animals carrying *ZNF251*WT or *ZNF251*KD3 tumors (50-60 mm^3^ were randomized into two groups (n=4/group), which were intraperitoneally administered vehicle or olaparib (10 mg/kg daily for four weeks). As expected, the volume and weight of *ZNF251*WT tumors were strongly reduced compared to those of their vehicle-treated counterparts (Fig. 1D, E). Remarkably, the tumor size and weight of the olaparib-treated *ZNF251*KD3 group were not reduced by olaparib treatment, which is consistent with the resistant phenotype (Fig. 1D, E). This shows that *ZNF251*KD breast cancer cells were resistant to olaparib treatment *in vivo*.

### ZNF251 haploinsufficiency confers resistance to multiple PARPis in BRCA1-mutated cells

To test whether the knockdown of *ZNF251* in breast cancer cells induces resistance to additional PARPis, we tested the resistance of *ZNF251*KD MDA-MB-436 clones to several potent PARPis, including niraparib (PARP1/2 inhibitor), veliparib (PARP1/2 inhibitor), NMS-P118 (selective PARP1 inhibitor), and stenoparib (PARP1/2 and PARP5a/5b inhibitor). Consistently, we observed that *ZNF251*KD breast cancer cells were resistant to all PARPis tested (Fig. 1F and Supplementary Fig. 4A).

To confirm our findings in a *BRCA1-*mutated ovarian cancer cell line, we knocked down *ZNF251* in the Ovcar8 cell line. Ovcar8 is a well-established human ovarian cancer cell line that has been shown to be methylated at the *BRCA1* promoter, resulting in decreased expression of *BRCA1* mRNA and protein(21). Subsequently, we tested the responses of these cells to various PARPis. We found that *ZNF251*KD ovarian cancer cells were also resistant to PARPis compared with *ZNF251*-wild type cells (Fig. 1G and Supplementary Fig. 4B). Importantly, in the absence of drug treatment, the growth rates of *ZNF251*KD breast and ovarian cancer cells were indistinguishable from those of their wild-type parental cells (Supplementary Fig. 5A, B).

To collect more evidence to support our findings, we also performed bioinformatic analysis of previously published PARPi resistance studies and found significantly lower expression of *ZNF251* in two olaparib-resistant breast cancer cell lines (MDA-MB-468 and SUM1315) when compared to their sensitive counterparts (Supplementary Fig. 6A)(12,13), showing that low expression of *ZNF251* is correlated with olaparib resistance. Furthermore, using CellMiner database analysis, we found that *ZNF251* expression was positively correlated with sensitivity to olaparib, cisplatin, and carboplatin in breast cancer cells (Supplementary Fig. 6B-D). Consistently, low *ZNF251* expression was correlated with worse survival in patients with breast cancer (22) (Supplementary Fig. 6E). Taken together, the downregulation of *ZNF251* was associated with resistance to olaparib and/or platinum derivatives in breast and/or ovarian *BRCA1*-mutated cancer cells. Moreover, cohorts of acute myeloid leukemia (AMLs) displayed low levels of *ZNF251* (Supplementary Fig. 6F), which may affect the outcome of clinical trials with PARPis(23).

### ZNF251 haploinsufficiency confers resistance to platinum-based drugs in BRCA1-mutated cells

Platinum-based anticancer drugs, including cisplatin, carboplatin, oxaliplatin, nedaplatin, and lobaplatin, are also commonly used as first-line chemotherapy regimens for cancer treatment. Mechanistically, these drugs form highly reactive platinum complexes that bind to and crosslink DNA in cancer cells. The mechanisms of action of platinum-based drugs and PARP are complementary in many ways and critically reliant on intracellular DNA damage(24). It has been previously reported that resistance to PARPis also results in platinum-based drug resistance(25,26). Therefore, we tested whether *ZNF251*KD breast cancer cells were resistant to platinum-based drugs. Experimentally, we treated *ZNF251*KD MDA-MB-436 clones with two platinum-based drugs, cisplatin and carboplatin, and tested their drug resistance. As expected, *ZNF251*KD MDA-MB-436 cells were resistant to both cisplatin and carboplatin (Fig. 1H, I). The IC_50_ of *ZNF251*KD breast cancer clones to cisplatin were 12.54-22.35 μM, whereas the IC_50_ for *ZNF251*WT cells was 1.93 μM. This indicated that *ZNF251* haploinsufficiency confers resistance to platinum-based drugs in *BRCA1*-mutated breast cancer cells. Interestingly, *ZNF251*KD *BRCA1*-mutated cells were not resistant to 5-fluorouracil (5-FU), which is primarily a thymidylate synthase (TS) inhibitor (data not shown).

### ZNF251 haploinsufficiency confers resistance to DNA polymerase theta (Polθ) inhibitors in BRCA1-mutated cells

Recent studies have suggested that HR-deficient cancer cells are sensitive to Polθ inhibitors owing to synthetic lethality(27,28). Moreover, HR-deficient PARPi-resistant cells are sensitive to DNA Polθ inhibitors(27,28). To test whether *BRCA1*-mutated *ZNF251*KD breast cancer cells are sensitive to DNA Polθ inhibitors, we treated MDA-MB-436 WT and *ZNF251*KD cells with the Polθ polymerase inhibitors ART-558 and ART-812(28) followed by a clonogenic assay. Interestingly, *BRCA1*-mutated *ZNF251*KD cells showed resistance to two Polθ inhibitors, ART812 and ART-558, compared to *BRCA1*-mutated *ZNF251*WT cells (Supplementary Fig. 7A, B). These results suggested that *ZNF251* haploinsufficiency confers resistance to Polθ and PARP inhibitors in *BRCA1*-mutated cells.

### ZNF251 haploinsufficiency induces upregulation of HR repair in BRCA1-mutated cells, resulting in olaparib resistance that can be reversed by a RAD51 inhibitor

To explore the molecular mechanisms by which *ZNF251*KD confers drug resistance to PARPi, we first examined DSBs by a neutral comet assay in *BRCA1*-mutated wild-type and *ZNF251*KD MDA-MB-436 cells treated with olaparib. We found that *ZNF251*KD MDA-MB-436 cells accumulated less olaparib-induced DSBs than their wild-type counterparts and were similar to those detected in *BRCA1*-restored cells (data not shown), which suggests a restoration of DSB repair in *ZNF251*KD cells. Reactivation of the HR pathway is a well-established mechanism associated with resistance to PARPis(21)(29). Therefore, we examined whether *ZNF251* haploinsufficiency affects DSB repair, especially the HR repair pathway. A specific reporter cassette measuring homologous recombination (HR) repair activity was used as previously described (10). Remarkably, we found that HR was dramatically activated only after olaparib treatment when compared to the *ZNF251* wild-type control in both breast cancer and ovarian cancer cells (Fig. 2A, B). To evaluate specific alterations in DSB repair HR pathways associated with *ZNF251* haploinsufficiency, we performed both RT-PCR and western blot analyses to examine the expression of the genes and proteins responsible for DSB repair, especially HR. Importantly, we observed a dramatic increase in the expression of RAD51 and RAD54, the key elements of the HR pathway, in olaparib-treated *ZNF251*KD MDA-MB-436 cells (Supplementary Fig. 8 and Fig. 2C, D), consistent with the stimulation of HR repair in olaparib-treated *ZNF251*KD cells (Fig. 2A, B). To confirm *ZNF251*KD induces PARPi resistance by restoring HR, we exposed *ZNF251*KD cells to RI-1, a potent and specific inhibitor of human RAD51(30). We found that treatment with RI-1 effectively inhibited HR repair (Fig. 2E) and reversed olaparib resistance in *ZNF251*KD cells (Fig. 2F), consistent with numerous previous observations that the restoration of HR repair can lead to PARPi resistance(16,17,29).

**Figure 2.**
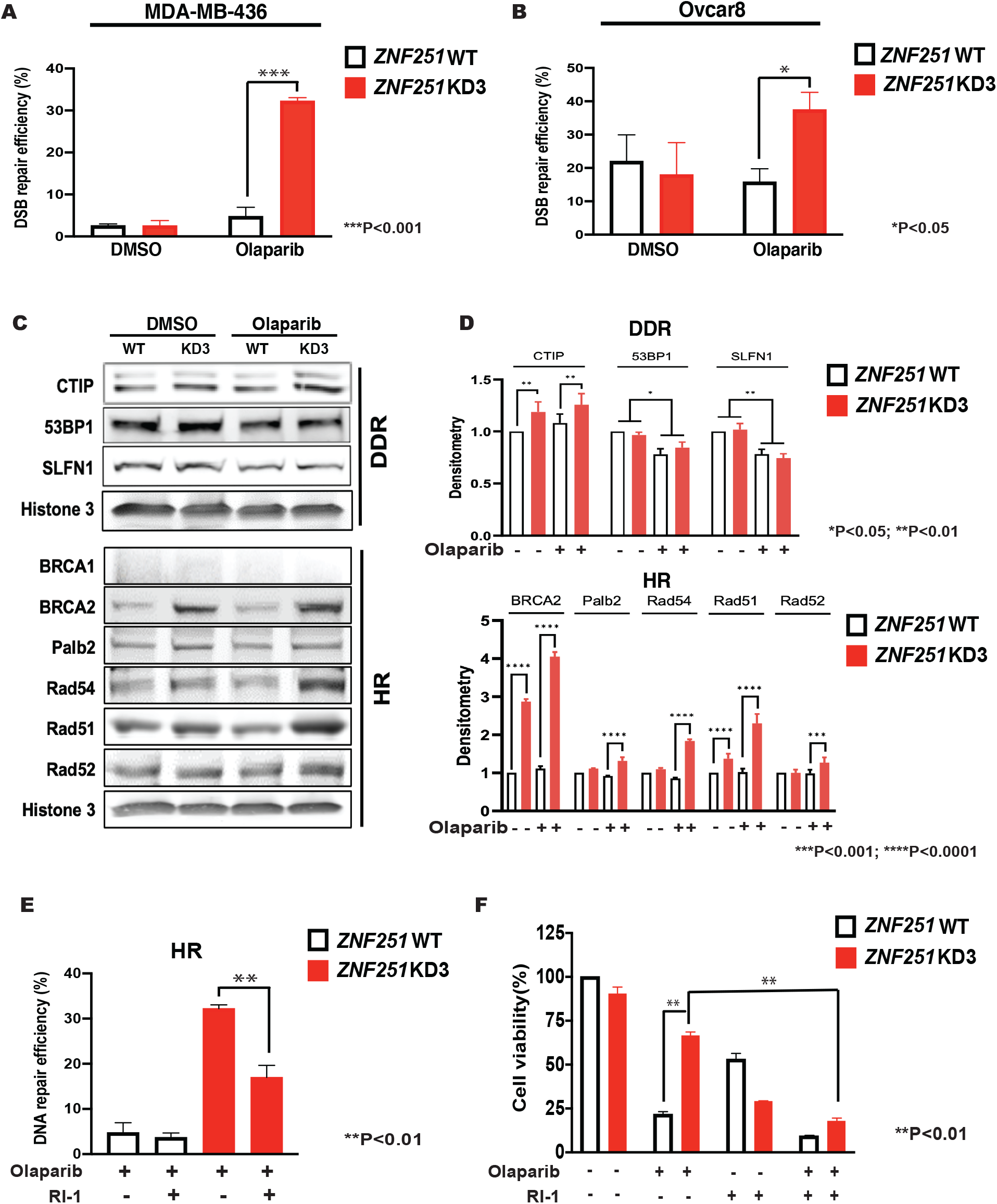
*ZNF251*KD resulted in upregulation of HR activity in olaparib-treated *BRCA1*-mutated cells. A, B. Reporter assay was carried out to determine the change of DSB repair pathways in both MDA-MB-436 (A) and Ovcar8 (B) cancer cells. C. Western blots were conducted to assess the expression levels of key components involved in double-strand break (DSB) repair, especially those related to homologous recombination (HR). D. Quantification of the western blot data. E, F. *ZNF251*WT and *ZNF251*KD MDA-MB-436 cells were treated with olaparib and RAD51 inhibitor RI-1 (16 μM) individually, as well as in combination for 3 days. Subsequently, a double-strand break (DSB) repair reporter assay was performed (E), followed by the assessment of olaparib resistance (F).

### ZNF251 haploinsufficiency causes replication fork protection in olaparib-treated BRCA1-mutated cells

It has been reported that resistance to PARPis-induced synthetic lethality may result from enhanced DSB repair as well as enhanced fork stabilization(31). To further explore the molecular mechanism underlying *ZNF251* haploinsufficiency-induced olaparib resistance, we tested whether *ZNF251*KD affected stabilization of the DNA replication fork. We performed a DNA replication fork protection assay (Supplementary Fig. 9A) in olaparib-treated wildtype and *ZNF251*KD cells. As expected, we found that olaparib treatment caused abundant DNA replication fork degradation in MDA-MB-436 wild-type cells, whereas *ZNF251*KD cells showed significant protection of the replication fork (Supplementary Fig. 9B). This result suggested that *ZNF251* haploinsufficiency protects replication forks from olaparib-induced degradation in *BRCA1*-mutated cells, leading to olaparib resistance. This observation is in line with multiple previous studies of PARPis resistance(32,33).

## Discussion

Four main mechanisms of acquired PARPi resistance have been identified in *BRCA1/2*-mutated cancer cells: alteration of drug availability, modulation of de-PARylation enzymes, restoration of HR, and enhanced replication fork stability(34,35). Using a positive whole-genome CRISPR/Cas9 library screen and several *BRCA1-*mutated breast and ovarian cancer cell lines, we discovered that haploinsufficiency of *ZNF251* which belongs to the Kruppel-associated box (KRAB) zinc-finger gene family cluster, caused resistance to multiple PARPis. Mechanistically, we discovered that *ZNF251* haploinsufficiency triggered PARPi resistance and was associated with the stimulation of HR repair, independent of the restoration of *BRCA1* expression.

The lack of BRCA1 is most likely compensated by the downregulation of 53BP1 (limits DNA end-resection) and upregulation of CtIP (promotes DNA end-resection) in olaparib-treated cells, causing an imbalance between CtIP and 53BP1, thus favoring DNA end-resection and generating substrates for HR (36). This conclusion is supported by the report that another zinc finger protein, ZNF432 played an important role in balancing the outcome of the PARPi response by modulating DNA resection(37).

In addition, *ZNF251* haploinsufficiency enhanced the expression of multiple key HR proteins, such as BRCA2, PALB2, RAD51, and RAD54, in olaparib-naïve and/or treated *BRCA1-*mutated cells. Similarly, ZNFs have been reported to regulate DSB repair(38,39). For example, ZNFs were capable of stimulating (ZNF506, ZNF384, and E4F1) and repressing (ZNF280C) NHEJ and HR(39-42).

Moreover, increased replication fork stability may also contribute to PARPi resistance in *ZNF251* haploinsufficient *BRCA1*-mutated cells. This speculation is supported by previous reports that replication fork stability confers PARPis resistance(25,33).

Although the detailed molecular mechanism of ZNF251 function in PARPi resistance still warrants further investigation, alterations in *ZNF251* expression may represent a novel diagnostic tool for prescreening patients with *BRCA1*-mutated tumors for potential treatment with RAD51 inhibitors.

## Authors’ Contributions

**Conception and design:** T. Skorski, J. Huang

**Development of methodology:** H. Li, S. Chatla, Z. Lian, T. Skorski, J. Huang

**Acquisition of data (provided animals, acquired and managed patients**,

**provided facilities, etc.):** H. Li, S. Chatla, X. Liu, Z. Tian, U. Vekariya, P. Wang, D. Kim, S. Octaviani, Z. Lian, D. Yang, H. Liu

**Analysis and interpretation of data (e.g**., **statistical analysis, biostatistics, computational analysis):** H. Li, S. Chatla, Z. Lian, X. Yu, KN. Chitrala, and J. Madzo **Writing, review, and/or revision of the manuscript:** H. Li, S. Chatla, J. Madzo, T. Skorski, and J. Huang.

**Administrative, technical, or material support (i.e**., **reporting or organizing Data, constructing databases):** H. Li, Z. Lian, G. Morton, Z. Feng, W. Childers, T. Skorski, and J. Huang

**Study supervision:** T. Skorski, J. Huang

## Acknowledgments

We sincerely thank Dr. Peter S. Klein at the University of Pennsylvania; Dr. Zhenkun Lou at Mayo Clinic; and Drs. Jean-Pierre Issa, Jaroslav Jelinek, Shumei Song, and Xiaoxin Chen at the Coriell Institute for Medical Research for their insightful comments and discussions. We thank all the members of the Skoriski lab and Huang lab for their help and discussions. We especially thank Steven Schneible at the Coriell Institute for assistance with manuscript editing.

## References

1. Barkauskaite E, Jankevicius G, Ahel I. Structures and Mechanisms of Enzymes Employed in the Synthesis and Degradation of PARP-Dependent Protein ADP-Ribosylation. Mol Cell 2015;58:935–46

2. Harrision D, Gravells P, Thompson R, Bryant HE. Poly(ADP-Ribose) Glycohydrolase (PARG) vs. Poly(ADP-Ribose) Polymerase (PARP) - Function in Genome Maintenance and Relevance of Inhibitors for Anti-cancer Therapy. Front Mol Biosci 2020;7:191

3. Rose M, Burgess JT, O’Byrne K, Richard DJ, Bolderson E. PARP Inhibitors: Clinical Relevance, Mechanisms of Action and Tumor Resistance. Front Cell Dev Biol 2020;8:564601

4. Gibson BA, Kraus WL. New insights into the molecular and cellular functions of poly(ADP-ribose) and PARPs. Nat Rev Mol Cell Biol 2012;13:411–24

5. Kim MY, Zhang T, Kraus WL. Poly(ADP-ribosyl)ation by PARP-1: ‘PAR-laying’ NAD+ into a nuclear signal. Genes Dev 2005;19:1951–67

6. Huang D, Kraus WL. The expanding universe of PARP1-mediated molecular and therapeutic mechanisms. Mol Cell 2022;82:2315–34

7. Hou P, Wu C, Wang Y, Qi R, Bhavanasi D, Zuo Z, et al. A Genome-Wide CRISPR Screen Identifies Genes Critical for Resistance to FLT3 Inhibitor AC220. Cancer Res 2017;77:4402–13

8. Tzavelas C, Bildirici L, Rickwood D. Production of stably transfected cell lines using immunoporation. Biotechniques 2004;37:276-8, 80-1

9. Guo C, Fordjour FK, Tsai SJ, Morrell JC, Gould SJ. Choice of selectable marker affects recombinant protein expression in cells and exosomes. J Biol Chem 2021;297:100838

10. Maifrede S, Le BV, Nieborowska-Skorska M, Golovine K, Sullivan-Reed K, Dunuwille WMB, et al. TET2 and DNMT3A Mutations Exert Divergent Effects on DNA Repair and Sensitivity of Leukemia Cells to PARP Inhibitors. Cancer Res 2021;81:5089–101

11. Hajji N, Garcia-Revilla J, Soto MS, Perryman R, Symington J, Quarles CC, et al. Arginine deprivation alters microglial polarity and synergizes with radiation to eradicate non-arginine-auxotrophic glioblastoma tumors. J Clin Invest 2022;132

12. Barrett T, Suzek TO, Troup DB, Wilhite SE, Ngau WC, Ledoux P, et al. NCBI GEO: mining millions of expression profiles--database and tools. Nucleic Acids Res 2005;33:D562–6

13. Gajan A, Sarma A, Kim S, Gurdziel K, Wu GS, Shekhar MP. Analysis of Adaptive Olaparib Resistance Effects on Cisplatin Sensitivity in Triple Negative Breast Cancer Cells. Front Oncol 2021;11:694793

14. Elstrodt F, Hollestelle A, Nagel JH, Gorin M, Wasielewski M, van den Ouweland A, et al. BRCA1 mutation analysis of 41 human breast cancer cell lines reveals three new deleterious mutants. Cancer Res 2006;66:41–5

15. Shalem O, Sanjana NE, Hartenian E, Shi X, Scott DA, Mikkelson T, et al. Genome-scale CRISPR-Cas9 knockout screening in human cells. Science 2014;343:84–7

16. Jaspers JE, Kersbergen A, Boon U, Sol W, van Deemter L, Zander SA, et al. Loss of 53BP1 causes PARP inhibitor resistance in Brca1-mutated mouse mammary tumors. Cancer Discov 2013;3:68–81

17. Dev H, Chiang TW, Lescale C, de Krijger I, Martin AG, Pilger D, et al. Shieldin complex promotes DNA end-joining and counters homologous recombination in BRCA1-null cells. Nat Cell Biol 2018;20:954–65

18. He YJ, Meghani K, Caron MC, Yang C, Ronato DA, Bian J, et al. DYNLL1 binds to MRE11 to limit DNA end resection in BRCA1-deficient cells. Nature 2018;563:522–6

19. Becker JR, Cuella-Martin R, Barazas M, Liu R, Oliveira C, Oliver AW, et al. The ASCIZ-DYNLL1 axis promotes 53BP1-dependent non-homologous end joining and PARP inhibitor sensitivity. Nat Commun 2018;9:5406

20. Yamaguchi H, Du Y, Nakai K, Ding M, Chang SS, Hsu JL, et al. EZH2 contributes to the response to PARP inhibitors through its PARP-mediated poly-ADP ribosylation in breast cancer. Oncogene 2018;37:208–17

21. Stordal B, Timms K, Farrelly A, Gallagher D, Busschots S, Renaud M, et al. BRCA1/2 mutation analysis in 41 ovarian cell lines reveals only one functionally deleterious BRCA1 mutation. Mol Oncol 2013;7:567–79

22. Gyorffy B. Survival analysis across the entire transcriptome identifies biomarkers with the highest prognostic power in breast cancer. Comput Struct Biotechnol J 2021;19:4101–9

23. Le BV, Podszywalow-Bartnicka P, Piwocka K, Skorski T. Pre-Existing and Acquired Resistance to PARP Inhibitor-Induced Synthetic Lethality. Cancers (Basel) 2022;14

24. Dilruba S, Kalayda GV. Platinum-based drugs: past, present and future. Cancer Chemother Pharmacol 2016;77:1103–24

25. Li H, Liu ZY, Wu N, Chen YC, Cheng Q, Wang J. PARP inhibitor resistance: the underlying mechanisms and clinical implications. Mol Cancer 2020;19:107

26. McMullen M, Karakasis K, Madariaga A, Oza AM. Overcoming Platinum and PARP-Inhibitor Resistance in Ovarian Cancer. Cancers (Basel) 2020;12

27. Zhou J, Gelot C, Pantelidou C, Li A, Yucel H, Davis RE, et al. A first-in-class Polymerase Theta Inhibitor selectively targets Homologous-Recombination-Deficient Tumors. Nat Cancer 2021;2:598–610

28. Zatreanu D, Robinson HMR, Alkhatib O, Boursier M, Finch H, Geo L, et al. Poltheta inhibitors elicit BRCA-gene synthetic lethality and target PARP inhibitor resistance. Nat Commun 2021;12:3636

29. Noordermeer SM, van Attikum H. PARP Inhibitor Resistance: A Tug-of-War in BRCA-Mutated Cells. Trends Cell Biol 2019;29:820–34

30. Budke B, Logan HL, Kalin JH, Zelivianskaia AS, Cameron McGuire W, Miller LL, et al. RI-1: a chemical inhibitor of RAD51 that disrupts homologous recombination in human cells. Nucleic Acids Res 2012;40:7347–57

31. Liao H, Ji F, Helleday T, Ying S. Mechanisms for stalled replication fork stabilization: new targets for synthetic lethality strategies in cancer treatments. EMBO Rep 2018;19

32. Rondinelli B, Gogola E, Yucel H, Duarte AA, van de Ven M, van der Sluijs R, et al. EZH2 promotes degradation of stalled replication forks by recruiting MUS81 through histone H3 trimethylation. Nat Cell Biol 2017;19:1371–8

33. Ray Chaudhuri A, Callen E, Ding X, Gogola E, Duarte AA, Lee JE, et al. Replication fork stability confers chemoresistance in BRCA-deficient cells. Nature 2016;535:382–7

34. Dias MP, Moser SC, Ganesan S, Jonkers J. Understanding and overcoming resistance to PARP inhibitors in cancer therapy. Nat Rev Clin Oncol 2021;18:773–91

35. Lord CJ, Ashworth A. Mechanisms of resistance to therapies targeting BRCA-mutant cancers. Nat Med 2013;19:1381–8

36. Cejka P, Symington LS. DNA End Resection: Mechanism and Control. Annu Rev Genet 2021;55:285–307

37. O’Sullivan J, Kothari C, Caron MC, Gagne JP, Jin Z, Nonfoux L, et al. ZNF432 stimulates PARylation and inhibits DNA resection to balance PARPi sensitivity and resistance. Nucleic Acids Res 2023;51:11056–79

38. Vilas CK, Emery LE, Denchi EL, Miller KM. Caught with One’s Zinc Fingers in the Genome Integrity Cookie Jar. Trends Genet 2018;34:313–25

39. Singh JK, Smith R, Rother MB, de Groot AJL, Wiegant WW, Vreeken K, et al. Zinc finger protein ZNF384 is an adaptor of Ku to DNA during classical non-homologous end-joining. Nat Commun 2021;12:6560

40. Moison C, Chagraoui J, Caron MC, Gagne JP, Coulombe Y, Poirier GG, et al. Zinc finger protein E4F1 cooperates with PARP-1 and BRG1 to promote DNA double-strand break repair. Proc Natl Acad Sci U S A 2021;118

41. Nowsheen S, Aziz K, Luo K, Deng M, Qin B, Yuan J, et al. ZNF506-dependent positive feedback loop regulates H2AX signaling after DNA damage. Nat Commun 2018;9:2736

42. Moquin DM, Genois MM, Zhang JM, Ouyang J, Yadav T, Buisson R, et al. Localized protein biotinylation at DNA damage sites identifies ZPET, a repressor of homologous recombination. Genes Dev 2019;33:75–89

