## Supplementary Fig. 1-9 and supplementary table 1 for "*ZNF251* haploinsufficiency confers PARP inhibitors resistance in *BRCA1*-mutated cancer cells through activation of homologous recombination"

### Supplementary Figure S1. A genome-wide CRISPR screen identified *ZNF251* whose haploinsufficiency conferred resistance to olaparib and is associated with *BRCA1* mutations.

**A**

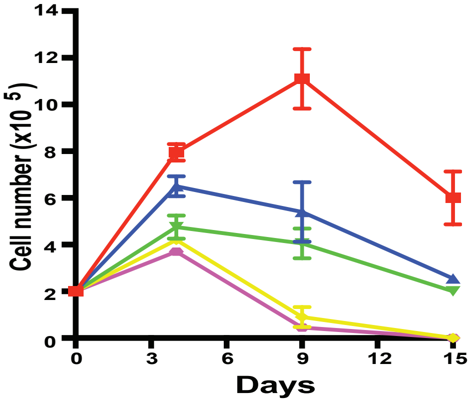


**B**

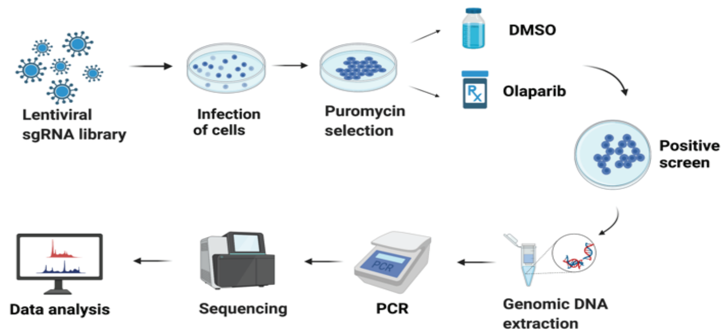


**C**

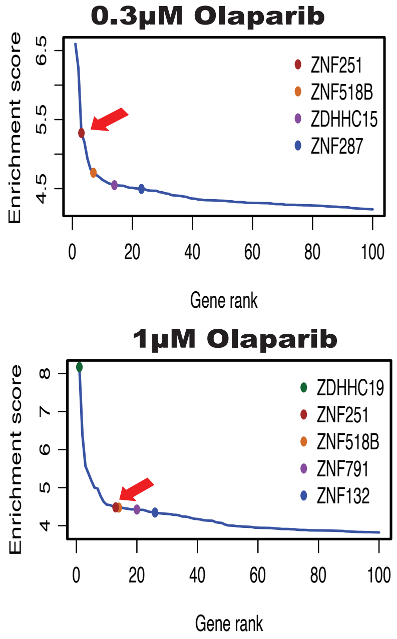


**D**

**0.3μM Olaparib**

| Genes | gRNA # recovered | Enrichment score |
|---|---|---|
| ***ZNF251*** | 5 | 5.30601517563597 |
| ***ZNF518B*** | 5 | 4.73026907245185 |
| ***TP53BP1*** | 5 | 2.60927595806458 |
| ***C20orf196*** | 4 | 3.36703503605691 |
| ***FAM35A*** | 6 | 2.99061242150602 |
| ***DYNLL1*** | 3 | 2.71699089440494 |
| ***EZH2*** | 4 | 2.54582918671737 |

**1μM Olaparib**

| Genes | gRNA # recovered | Enrichment score |
|---|---|---|
| ***ZNF251*** | 6 | 4.48178570537391 |
| ***ZNF518B*** | 6 | 4.47704228454464 |
| ***TP53BP1*** | 6 | 1.73363976845943 |
| ***C20orf196*** | 5 | 2.61559411937482 |
| ***FAM35A*** | 6 | 2.60467560322601 |
| ***DYNLL1*** | 5 | 2.12267271882414 |
| ***EZH2*** | 6 | 2.13750352374994 |

**E**

***ZNF251*** **WT1** CCTTCCAGGATGTGGCCG----TG-------TACTTCTCTCAGGCGGTGGGGCTTAAGAG
***ZNF251*** **KD1** CCTTCCAGGATGTGGCCGGCGCTGAACGAACCACTTCTCTCAGGCGGTGGGGCTTAAGAG

***ZNF251*** **WT2** GGTCCTGAATCTTCTGGGAGCTGAGGAACCAGATATCTTGAAAAGCTGCCAGAAAGGTGA
***ZNF251*** **KD2** GGTCCTGAATCTTCTGGGCACTGAGGAACCAGATATCTTG-AAAGCTGCCAGAAAGGTGA

***ZNF251*** **WT3** GGCCGTGAGGGCAAACTCAAAGAGCGCGTGGGAAATTCTGCCGGGCAGAGTTTGAACAAA
***ZNF251*** **KD3** GGCCGTGAGGGCAAACTCAAAGA--GCGTGGGAAATTCTGCCGGGCAGAGTTTGAACAAA

**F**

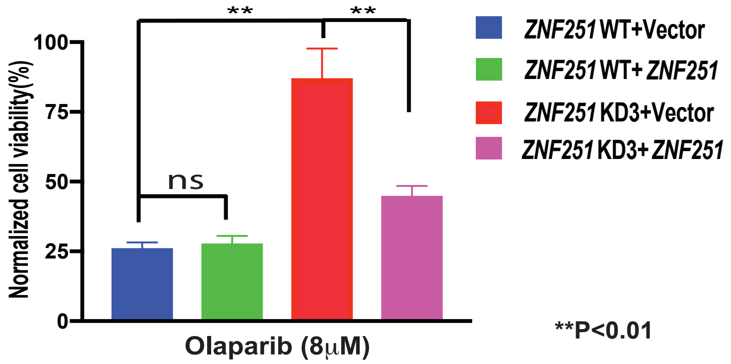


Supplementary Fig. S1. A genome-wide CRISPR screen identified *ZNF251* whose haploinsufficiency conferred resistance to olaparib and is associated with *BRCA1* mutations. A. The growth curve of MDA-MB-436 breast cancer cells treated with various doses of olaparib. B. A simplified scheme of the olaparib resisatance CRISPR screen with MDA-MB-436 cells. C.Enrichment of specific sgRNAs that target each gene after 14 days of olaparib treatment and identification of top candidate genes. The x axis represents enriched genes, and the y axis represents sgRNA enrichment score, which was calculated using (sgRNA number from the reads)/(sgRNA number in the library)/ log2 (average abundance). Arrow indicates *ZNF251* gene. D. The gRNAs # recovered from our screen and the enrichment score for *ZNF251*, *ZNF518B*, and several known genes whose loss-of-function causes olaparib resistance were listed. E. Sanger sequencing data of three *ZNF251*KD clones. F. *ZNF251*WT or *ZNF251*KD3 cells transfected with pcDNA3.1 vector or *ZNF251* cDNA on pcDNA3.1 vector were treated with 8μM olaparib for 72 hours and cell viability was measured and normalized to DMSO-treated corresponding cells. The results represent three independent experiments.

**Supplementary Figure S2. *ZNF251*KD resulted in increased proliferation and decreased apoptosis of breast cancer cells.**

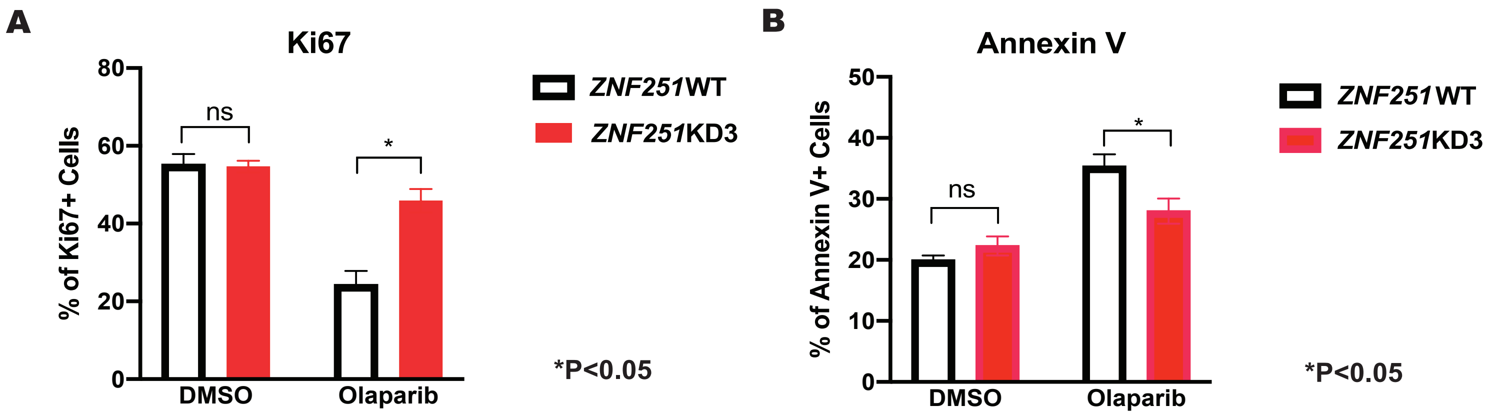


Supplementary Fig. S2. *ZNF251*KD resulted in increased proliferation and decreased apoptosis of breast cancer cells. A. Ki67 staining was conducted on WT and *ZNF251*KD3 breast cancer cells treated with either DMSO or olaparib. B. Annexin V staining was performed on WT and *ZNF251*KD3 breast cancer cells treated with either DMSO or olaparib.

**Supplementary Figure S3. The three individual clones with *ZNF251*KD displayed resistance to olaparib in the *BRCA1*- but not *BRCA1*+ background breast cancer cells.**

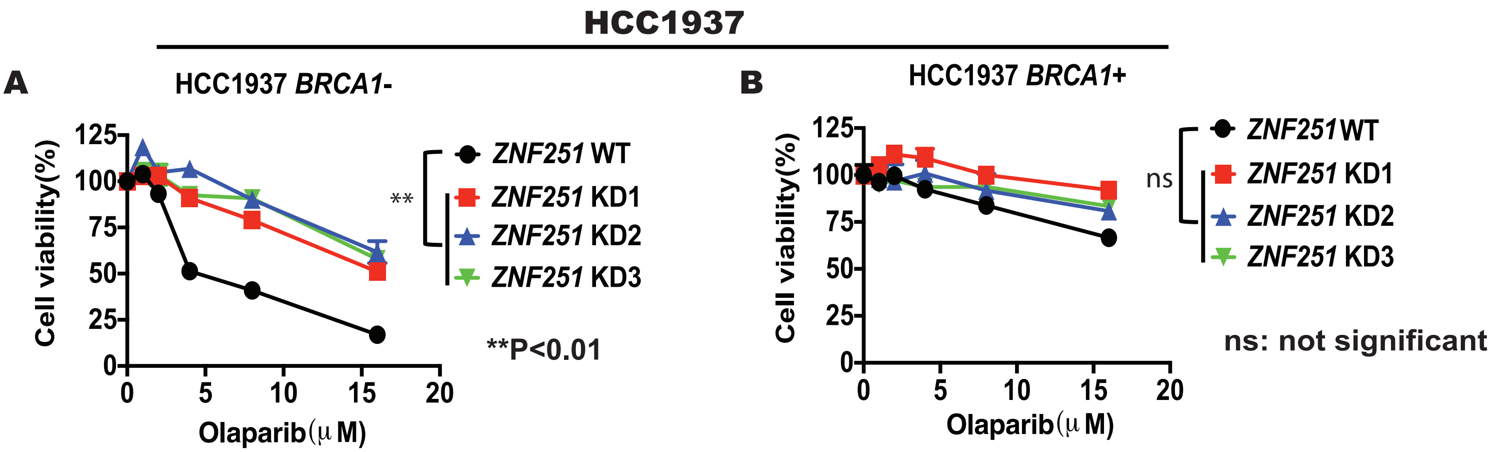


Supplementary Fig. S3 The three indivitual clones with *ZNF251*KD showed resistance to olaparib in *BRCA1*- but not *BRCA1*+ background breast cancer cells. A, B. *ZNF251*KD was constructed in HCC1937 (*BRCA1*- or *BRCA1*+) lines and the resistance to olaparib was measured. The results represent three independent experiments.

**Supplementary Figure S4. The three individual clones with *ZNF251*KD in both breast and ovarian cancer displayed resistance to the PARP inhibitor veliparib..**

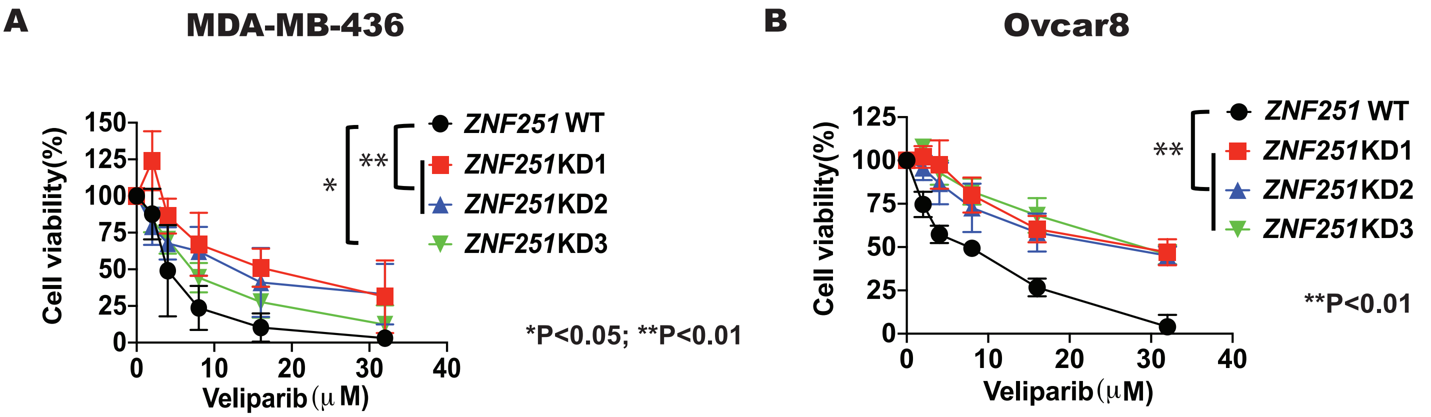


Supplementary Fig. S4 The three indivitual clones with *ZNF251*KD in both breast and  ovarian cancer displayed rsistance to the PARPi veliparib. A, B. Cell growth curve of WT and three ZNF251KD single clone breast (MDA-MB-436) and ovraian (Ovcar8) cancer cells following treatment with veliparib. The results represent three independent experiments.

**Supplementary Figure S5. The three individual clones with *ZNF251*KD in both breast and ovarian cancer showed no growth difference compared to the *ZNF251* wildtype cells in the absence of PARPi.**

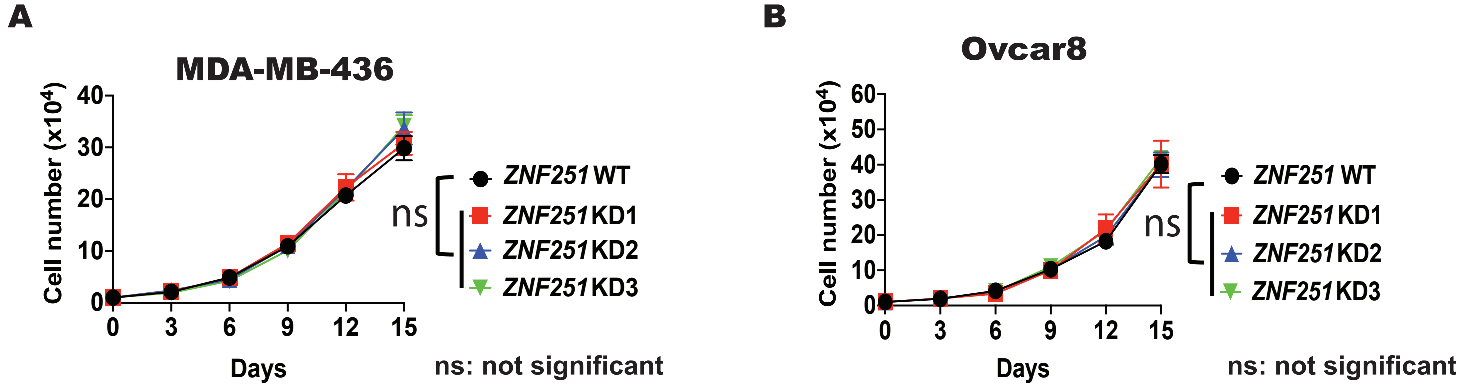


Supplementary Fig. S5 The three indivitual clones with *ZNF251*KD in both breast and ovarian cancer showed no growth difference compared to the *ZNF251*WT cells in the absence of PARPis. A, Cell growth was measured in *ZNF251*WT and three indivitual *ZNF251*KD MDA-MB-436 breast cancer cells. B. Cell growth was measured in *ZNF251*WT and three indivitual *ZNF251*KD Ovcar8 ovarian cancer cells.

### Supplementary Figure S6. Bioinformatic analysis showed that *ZNF251* expression is correlated with PARPi and platinum drugs sensitivity as well as prognosis of breast cancer patients

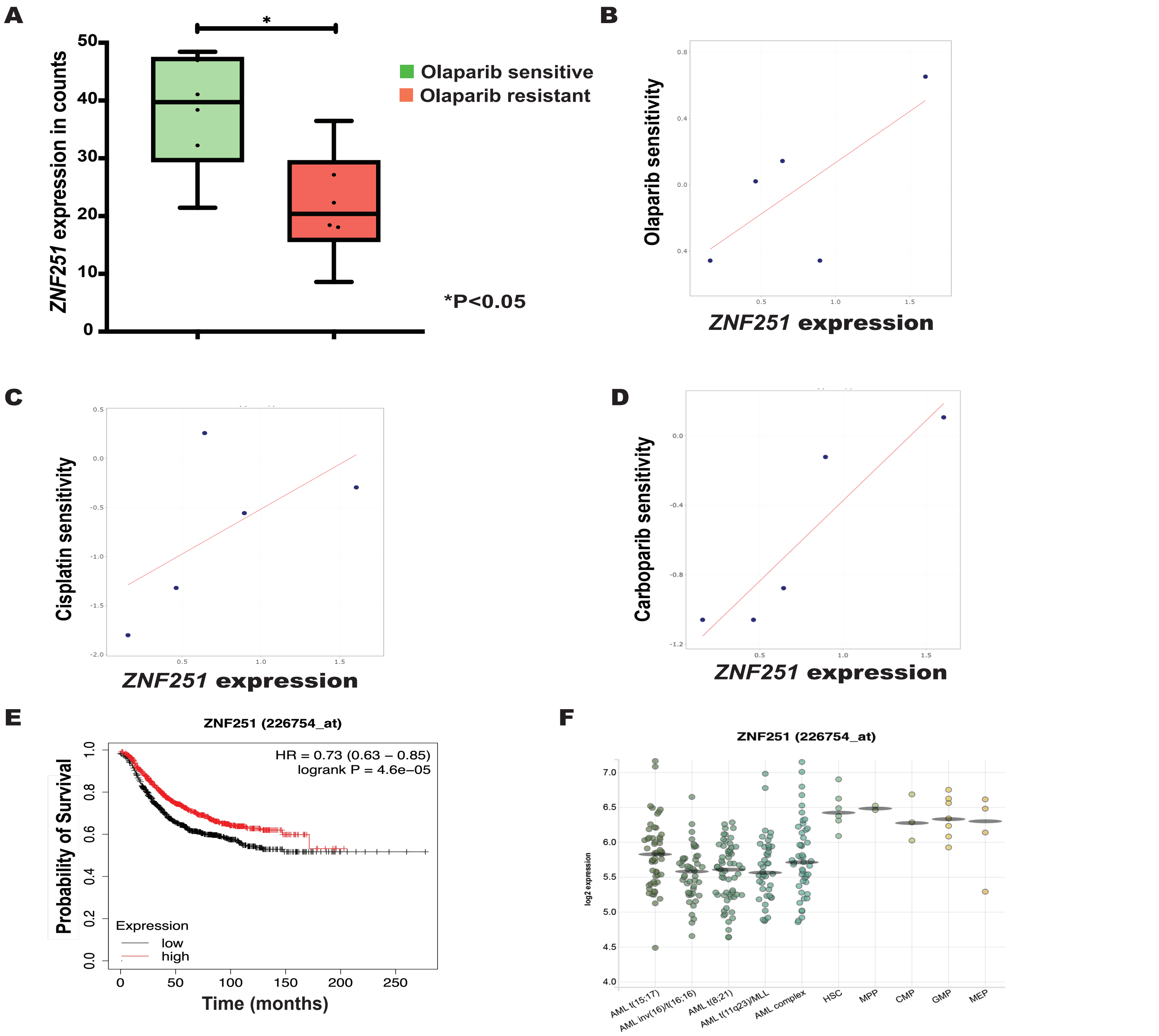


Supplementary Fig. S6. Bioinformatic analysis showed that *ZNF251* expression is corelated with PARPi and platinum drugs sensitivity as well as prognosis of breast cancer patients. A. *ZNF251* expression was correlated with olaparib resistance of breast cancer cells. Box-whisker plots indicating the *ZNF251* expression values of olaparib sensitive and resistant cells collected from the Gene Expression Omnibus (GEO) database datasets. Green color plot represents sensitivity towards PARPi whereas red color plot represents resistance towards PARPi. Box-whisker plot for olaparib sensitive and resistant cells was generated using the GEO dataset GSE165914. Y-axis represents the expression of *ZNF251* in the respective cells. Statistical analysis was performed using the 2-tailed Student's t test. *p-value<0.05. B-D. *ZNF251* expression is positively correlated with sensitivity to olaparib, cisplatin and carboplatin of the breast cancer cells by CellMiner database analysis. E. Low *ZNF251* expression is correlated with worse survival for breast cancer patients using online Kaplan-Meier plotter, which integrates gene expression and clinical data from 2,032 patients. F. Expression of ZNF251 in AMLs displaying the indicated karyotype and in normal hematopoietic cells (HSC = hematopoietic stem cells, MPP = multipotent progenitors, CMP = common myeloid progenitors, GMP = granulocyte-monocyte progenitors, MEP = megakaryocyte-erythrocyte progenitors) according to Bloodspot database. https://servers.binf.ku.dk/bloodspot/?gene=ZNF251&dataset=normal_human_v2_with_AMLs.

**Supplementary Figure S7. *ZNF251*KD confers resistance to Polθ inhibitors**

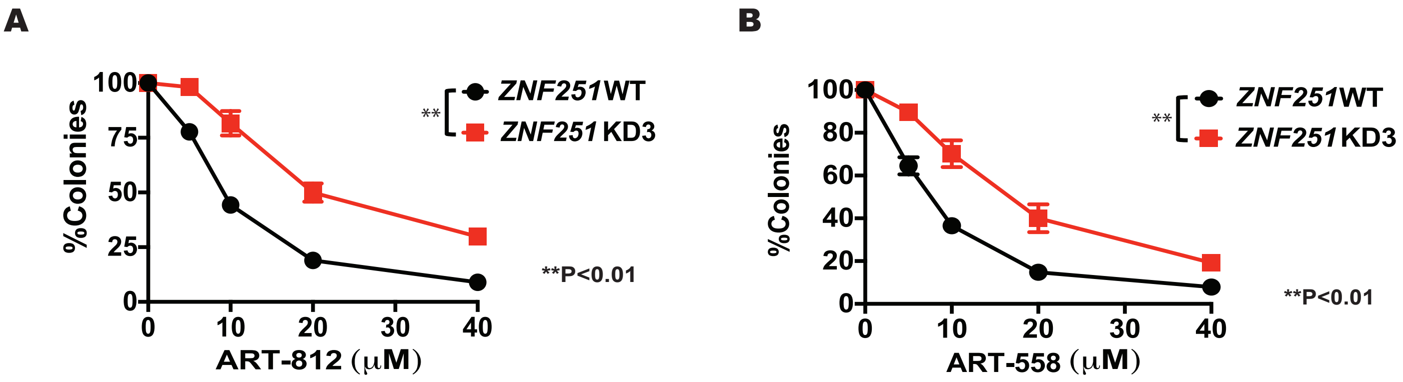


Supplementary Fig. S7. *ZNF251*KD confers resistance to Polθ inhibitors. A, B. Sensitivity of MDA-MB-436 *ZNF251*WT and MDA-MB-436 *ZNF251*KD cells to DNA polymerase θ inhibitors ART-812 and ART-558 at indicated concentrations. The results represent mean % colonies ± SDs when compared to DMSO-treated cells.

**Supplementary Figure 8.** Haploinsufficiency of *ZNF251* results in the upregulation of gene transcription for homologous recombination (HR) factors.

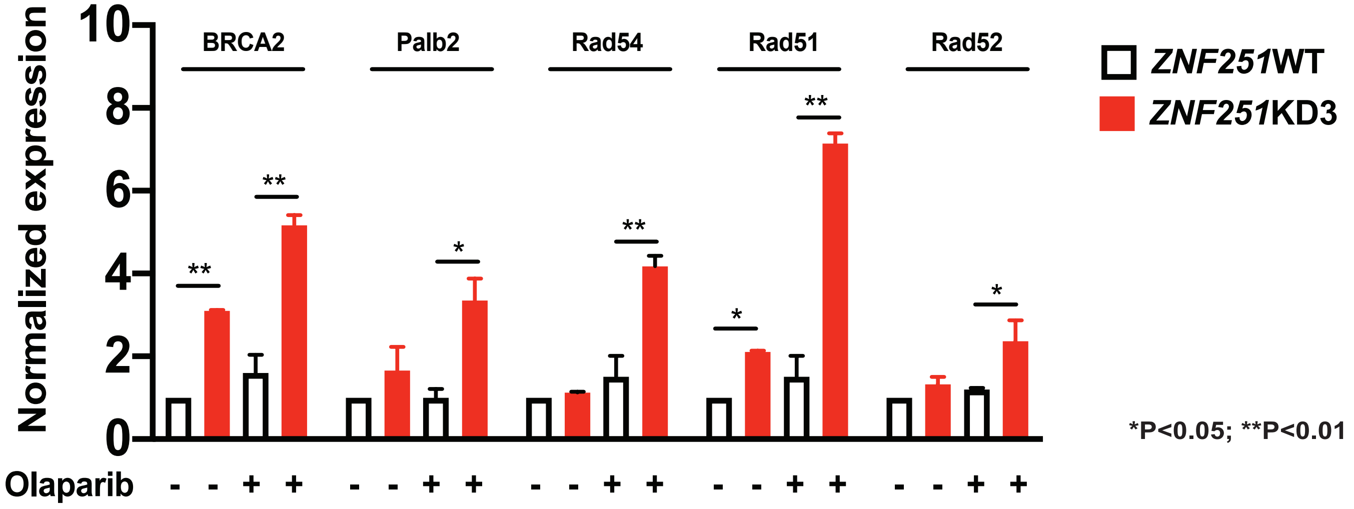


Supplementary Fig.8 Haploinsufficiency of *ZNF251* results in the upregulation of gene transcription for homologous recombination (HR) factors. The expression levels of *BRCA2*, *PALB2*, *RAD54*, *RAD51*, and *RAD52* were measured by RT-PCR in *ZNF251*WT and *ZNF251*KD3 breast cancer cells treated with either DMSO or olaparib.

**Supplementary Figure 9. *ZNF251* haploinsufficiency causes replication fork protection in olaparib-treated *BRCA1*-mutated cancer cells.**

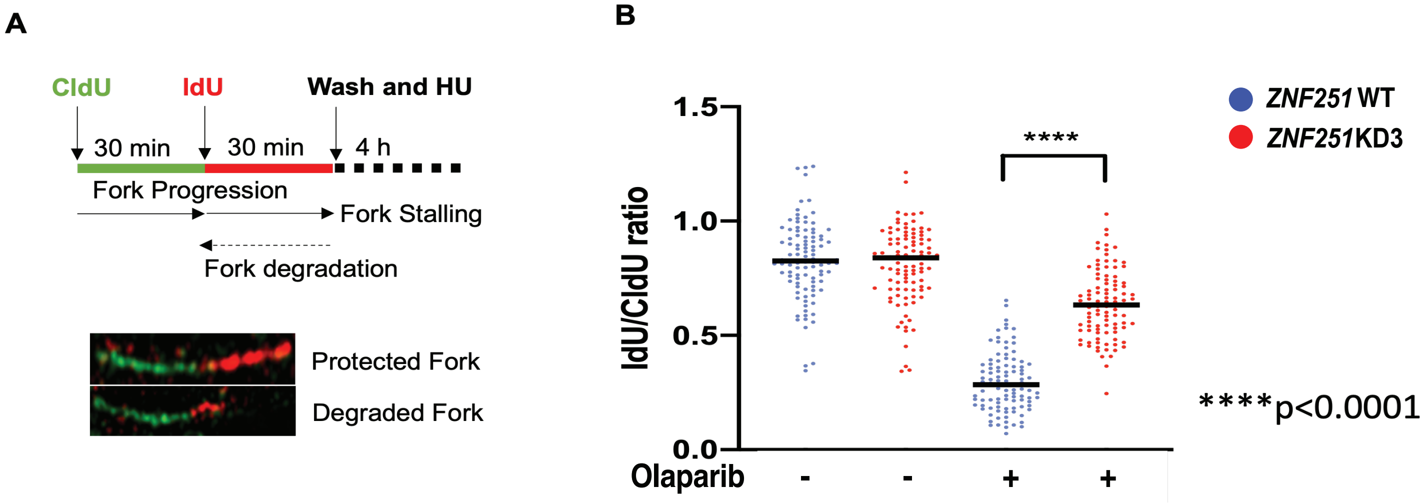


Supplementary Fig.9 *ZNF251* haploinsufficiency causes replication fork protection in olaparib-treated *BRCA1*-mutated cancer cells.A. Top: schematic representation of the protection of nascent DNA at stalled replication forks employing DNA fiber assay. Bottom: representative images of protected and degraded DNA fibers. B. Graph summarizing the quantification of IdU/CIdU ratio for n = 100 DNA fibers analyzed per sample for each experiment (Cells were treated 5 mM Olaparib ). The graph is representative of 2 independently performed experiments. Significance was calculated with the Mann–Whitney U-test, and bar indicated the median for each sample. ****P < 0.0001 differences between samples.

**Supplementary Table S1. List of primers used in this study**

***ZNF251*** **gRNA oligos**

gRNA1Forward: 5’-CACCGCGTGTACTTCTCTCAGGCGG (Exon 3)
gRNA1Reverse: 5’-AAACCCGCCTGAGAGAAGTACACGC

gRNA2Forward: 5’-CACCGGGGAGATCAACTCCGGCTTA(Exon 4)
gRNA2Reverse: 5’-AAACTAAGCCGGAGTTGATCTCCCC

gRNA3Forward: 5’-CACCGGGCAAACTCAAAGAGCGCGT (Exon 5)
gRNA3Reverse: 5’-AAACACGCGCTCTTTGAGTTTGCCC

**T7E1 primers**

gRNA1 Forward: 5’-CTGGGTCCCCGTGTAGCTGAAGTT
gRNA1 Reverse: 5’-CATCAGCTCTCCTGCCACTGGAGG

gRNA2 Forward: 5’-CTCCTTCAGTACTTAGTCTCAGCCC
gRNA2 Reverse: 5’-TGTGGTAGAAGCTCCTATAAACCATCA

gRNA3 Forward: 5’-ATTCTGAGGTTGGGACCAAGAAGG
gRNA3 Reverse: 5’-CACCGGCCACATTCGTACGGCTTC

**RT-PCR primers**

*RAD51*
Forward Sequence: TCTCTGGCAGTGATGTCCTGGA
Reverse Sequence: TAAAGGGCGGTGGCACTGTCTA

*RAD52*
Forward Sequence: GGCAGTTCAACATCAAGCCCTTG
Reverse Sequence: CTTCGGAAGTGGCACCATGCTA

*RAD54*
Forward Sequence: ACGGCGTTAGTGGTTTGTCCTC
Reverse Sequence: GCAGCATGTAGCTTCTCTCCTG

*BRCA2*
Forward Sequence: GGCTTCAAAAAGCACTCCAGATG
Reverse Sequence: GGATTCTGTATCTCTTGACGTTCC

*PALB2*
Forward Sequence: GGAGCTGCATAAACATTCCGTCG
Reverse Sequence: CTACGGAACAGGAACCTGAAGG
